# mRNA challenge predicts brain cancer immunogenicity and response to checkpoint inhibitors

**DOI:** 10.1101/2023.03.18.532056

**Authors:** Paul Castillo, Elizabeth Ogando-Rivas, Hilary Geffrard, Alfonso Pepe, Ruixuan Liu, Duy T Nguyen, Diego I Pedro, Dingpeng Zhang, Anna DeVries, Sadeem Qdaisat, Aida Karachi, Maryam Rahman, Frances Weidert, Rowan Milner, Jianping Huang, Natalie L. Silver, John Ligon, Derek Li, Ji-Hyun Lee, Sheila Carrera-Justiz, Duane A Mitchell, Hector Mendez-Gomez, W Gregory Sawyer, Elias J Sayour

**Affiliations:** University of Florida Department of Pediatrics, Division of Pediatric Hematology Oncology, Gainesville, FL, 32610; University of Florida Lillian S. Wells Department of Neurosurgery, Preston A. Wells, Jr. Center for Brain Tumor Therapy, Gainesville, FL, 32610, USA; University of Florida Department of Mechanical and Aerospace Engineering, Gainesville, FL, 32610; University of Florida, College of Veterinary Medicine, Gainesville FL, 32610; Cleveland Clinic, Center of Immunotherapy and Precision Immuno-Oncology/Head and Neck Institute, Cleveland, OH 44106; University of Florida, Department of Biostatistics, Gainesville, FL 32610

**Author notes:** contributed equally (Paul Castillo, Elizabeth Ogando-Rivas, Hilary Geffrard, Alfonso Pepe). Corresponding author: Elias Sayour, M.D., Ph.D. University of Florida Lillian S. Wells Department of Neurosurgery, Preston A. Wells, Jr. Center for Brain Tumor Therapy, Gainesville, FL, 32610, USA.

**Keywords:** mRNA, brain cancer immunogenicity, response to immune checkpoint inhibitors

## Abstract

To prospectively determine whether brain tumors will respond to immune checkpoint inhibitors (ICIs), we developed a novel mRNA vaccine as a viral mimic to elucidate cytokine release from brain cancer cells in vitro. Our results indicate that cytokine signatures following mRNA challenge differ substantially from ICI responsive versus non-responsive murine tumors. These findings allow for creation of a diagnostic assay to quickly assess brain tumor immunogenicity, allowing for informed treatment with ICI or lack thereof in poorly immunogenic settings.

---

In patients with primary glioblastoma (GBM), we demonstrated that increased tumor infiltrating lymphocytes (TILs) correlate with improved survival^1^, paving the way for immunotherapeutic strategies to deplete Tregs or promote effector TILs in the tumor microenvironment (TME). We have advanced these findings to prospectively determine whether a given brain tumor will be expected to have T cell infiltration and respond to immune checkpoint inhibitors (ICIs). We hypothesize that immune recruitment is dictated by cancer cell expression of pathogen recognition receptors (PRRs) that activate in response to oncogenic stress from DNA/RNA released from apoptotic cells.

To prospectively determine whether individual brain tumors respond to ICIs, we developed a novel mRNA formulation composed of multilamellar mRNA aggregates^2^ (**Fig. 1a**) as a broad screen to determine cancer cell immunogenicity. These RNA-lipid particles (LPs) activate intracellular PRRs^2^ but can also stimulate MYD88 dependent sensors (**Fig. 1b**), such as endosomal toll-like receptors, providing a single approach to simultaneously activate multiple PRRs. We challenged murine brain tumor lines known to respond or resist treatment following immune checkpoint inhibitors (ICIs) with these RNA-LPs and analyzed downstream production of proinflammatory cytokines as predictors of ICI response (**Fig. 1c**). Following transfection, we expected that murine tumors responsive to ICIs would have a distinct cytokine response signature, while unresponsive tumors would not elicit production of proinflmmatory cytokines. We chose brain tumor lines GL261 and SMA-560 as ICI responsive and KR158b-luc and CT-2A as ICI unresponsive based on previous literature^3,4^ and our experience (**Fig. 1d**). In 2D in vitro culture, we successfully transfected these cell lines with GFP mRNA (**Fig. 1e**). Following mRNA challenge, ICI responsive tumors GL261 and SMA-560 showed increases in pro-inflammatory (**Fig. 1f**) suggesting these (as expected) to be the most immunogenic tumors.

**Fig. 1:**
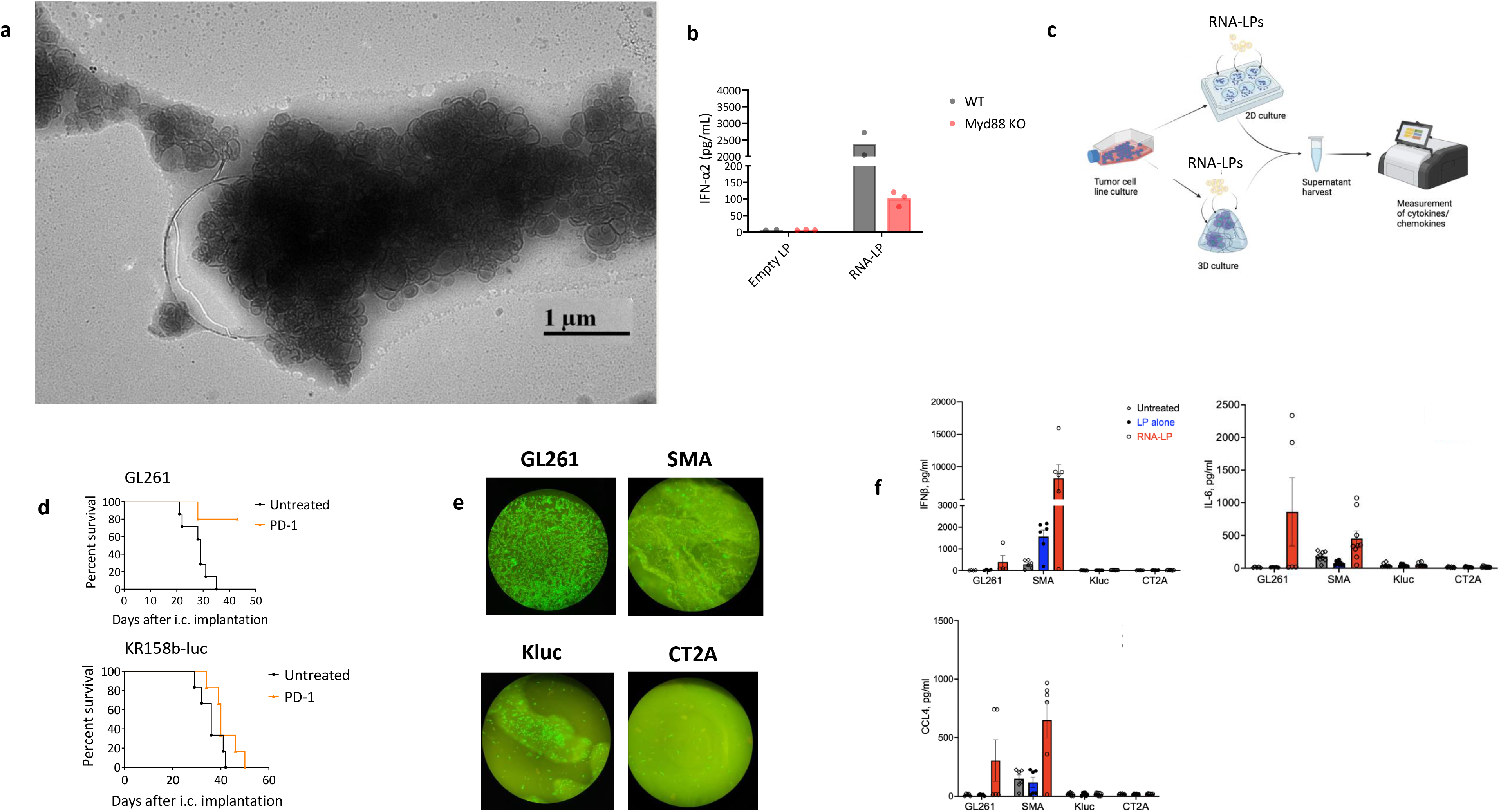
Cytokine response to mRNA challenge predicts response to anti-PD-1 in murine brain cancer lines. **a,** Sheets of RNA-LP aggregates by cryo-electron microscopy, **b,** Wild-type C57bl/6 mice or MYD88 knock-out (KO) mice bearing pulmonary B16F10-OVA tumors were intravenously infused with OVA specific RNA-LPs and serum collected 6h after infusion for assessment of IFN-α2 by ELISA. **c,** Study schema. **d,** (top) GL261 or (bottom) KR158b-luc (K-luc) tumors were implanted intracranially and treated i.p. with PD-1 monoclonal antibodies as loading dose (first day), and administered every 3-4 days thereafter. **e,** GFP transfection efficiency of brain tumor cell lines in vitro. **f,** Cytokine analysis of transfected cell supernatants. Images created with Biorender.com.

Since cell lines have high failure rate, and to further develop this work into a commercially viable tool to predict ICI response, we developed 3D modeling of glial tumors for mRNA challenge and immunogenicity prediction. We enrolled canines with primary gliomas onto clinical trials as previously described^2^, and began growing their tumoroids in 3D using liquid-like solid (LLS) technology that rapidly allows for simultaneous growth of 96 patient-derived microtumors as spheroids (**Fig. 2a**). ^5,6^ In this system, we show that canine gliomas form spheroids with LLS serving as the scaffold (**Fig. 2b**). In separate studies from murine lines using the same pipeline as in **Fig. 1c**, we successfully perfused 3D murine tumoroids with RNA-LP as evidenced by GFP expression (**Fig. 2c**) demonstrating ability to set up real-time 3D assays for mRNA challenge to predict immunogenicity (**Fig. 2d**). These data provide proof of concept that we may be able to prospectively enroll patients and grow personalized tumoroids for mRNA challenge. To demonstrate that mRNA perfusion of 3D tumoroids would elicit similar cytokine response observed with 2D culture of cell-lines, we challenged 3D tumoroids with GFP mRNA and elicited a similar pattern of increased IFN-β, IL-6 and CCL4 (**Fig 2e**) in GL261 and SMA-560 but not in KR158b-luc or CT-2A. Thus, 3D modeling of murine gliomas confirmed cytokine response signature unique to checkpoint responsive malignancies.

**Fig. 2:**
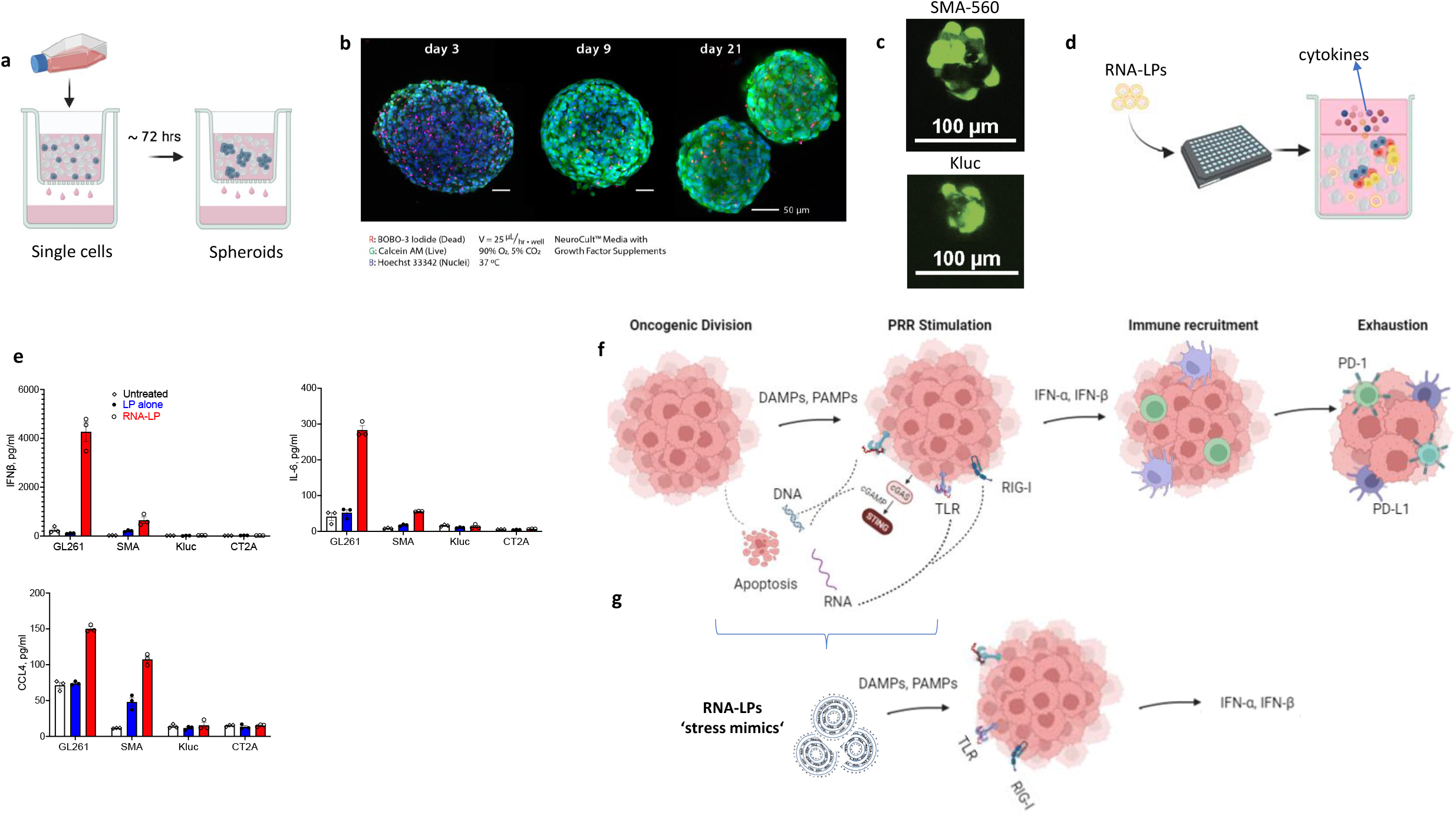
Rapid 3D modeling enables detection of cytokine response to mRNA challenge. **a,** Schematic of tumors grown in 3D with LLS. **b,** Longitudinal imaging of primary canine glioma grown in 3D using LLS. **c,** GFP expression following mRNA transduction of 3D tumoroids. **d,** Schematic of RNA-LP challenge of tumoroids in 3D LLS. **e,** Cytokine analysis from liquid culture media of transfected 3D tumoroid cell supernatants. **f, g,** Proposed mechanism for induction of cancer immunogenicity (f) and use of RNA-LP as stress mimic to predict immunogenicity (**g**). Images created with Biorender.com.

Cancer cells can evolve to subvert immune recognition while mediating recruitment of myeloid derived suppressor cells and tumor associated macrophages that exclude T cells from the TME.^7^ Alternatively, immunogenic tumors can alert DCs and T cells through innate production of cytokines/chemokines eliciting adaptive immunity that is stymied through expression of immune checkpoints.^8^ These innate cytokines (e.g., IFN-α) are produced early on during tumorigenesis from checkpoint sensitive tumors in response to nucleic acid (DNA, RNA) release elucidating the development and regulation of our immune system’s innate response. Rapid cell division creates competition and stress among cancer cell subpopulations, leading to apoptosis and release of nucleic acid or pathogen associated molecular patterns (PAMPs) alongside damage associated molecular patterns (DAMPs). These PAMPs can trigger PRRs in the TME inducing type I interferon responses that recruit DCs/T cells predisposing cancer immunogenicity and exhaustion (**Fig. 2f**). Alternatively, tumor cells may evolve mechanisms or grow from stem progenitor states lacking PRR machinery that stymie this process to prevent T cell recruitment and infiltration. These ‘hot’ versus ‘cold’ tumors are difficult to rapidly predict. By encapsulating single/double-stranded stranded elements^2^ and eliciting DAMPs through cationic charge, we developed a multilamellar RNA-LP approach to rapidly predict cancer cell immunogenicity through induction of type I interferon responses as a surrogate for ICI responsiveness (**Fig 2g**). Our results indicate that cytokine signatures are significantly different under immunogenic stress from mRNA challenge and may predict ICI responsiveness. While a limitation of our study is that these cytokine signatures have thus far been observed in spheroids generated from murine tumor lines, we have demonstrated the technical feasibility of generating these models from real-time canine tumors and could incorporate this test as a meaningful correlative study in a prospective human clinical trial of ICIs for brain tumors to validate these findings. Recent findings have uncovered that DNA repair machinery (as opposed to neoantigen burden) is significantly implicated in poorly immunogenic tumors^9^ substantiating the premise that cancer immunogenicity can be triggered through release of DAMPs and PAMPs. Developing an assay to quickly assess how tumor cells respond to these triggers could be leveraged to prospectively manage patients. Our findings may allow for creation of a diagnostic assay to quickly assess whether some malignancies are immunogenic, allowing for informed treatment with checkpoint inhibitors or lack thereof in poorly immunogenic settings.

## Acknowledgements

The authors would like to thank Lana Fagman and Kaylee Young for veterinary trials assistance.

## Methods

### Cell lines and mice

Murine glioma cell lines (i.e., GL261, KR158b-luc, SMA-560, CT2A) were used. GL261 (Glioma 261) cells were supplied by the National Cancer Institute (NCI). B16F10-OVA and KR158b-luc (K-luc) gliomas were obtained as previously described.^2^ SMA-560 and CT2A cell lines were obtained from Sigma-Aldrich. GL261 cells were maintained in Dulbecco’s Modified Eagle Medium (DMEM) F12/Glutamax (Gibco 10565018, Waltham, MA) with 10% fetal bovine serum (FBS) and 1% penicillin/streptomycin. KR158-luc cells were maintained in DMEM without sodium pyruvate (Gibco 11965-092, Waltham, MA) with 10% FBS and 1% penicillin/streptomycin. SMA-560 cells were maintained in Minimum Essential Medium (MEM, Gibco 10373017, Waltham, MA) with 10% FBS and 1% penicillin/streptomycin. CT2A cells were maintained in DMEM with 10% FBS and 1% penicillin/streptomycin. Cell were incubated at 37C 5% CO2. Wild-type C57bl/6 mice and B6.129P2(SJL)-*Myd88^tm1.1Defr^*/J (MYD88 knock-out) mice were obtained from the Jackson Laboratories and cared by the University of Florida (UF) Animal Care Services team. All in vivo experiments were approved by the UF Institutional Animal Care and Use Committee (IACUC).

### mRNA and RNA-LP complexation

Synthetic mRNA encoding for full-length model antigen (i.e., CleanCap GFP, catalog L-7601 and CleanCap ovalbumin, OVA, catalog L-7610) were purchased from TriLink Biotechnologies (San Diego, CA). mRNA constructs were validated based on sizing gel-electrophoresis, and nanodrop quality/bioanalysis. RNA-LPs were generated as previously described.^2^ Briefly, DOTAP (1,2-dioleoyl-3-trimethylammonium-propane - chloride salt) particles that are positively charged were complexed with negatively charged synthetic mRNA at the RNA:LP ratio of 1:15 as previously described.^2^ After 15 minutes of complexation at room temperature, the resulting product was used for downstream experiments. For in vitro experiments, RNA-LPs were delivered in 45μL (5ug, fixed mRNA concentration of 1 μg/μL). For in vivo experiments, RNA-LPs were delivered as 25 μg doses per mouse in 200uL.

### Imaging of RNA-LPs

Complexed RNA-LPs were snap frozen in liquid nitrogen and then imaged by cryo transmission electron microscopy as previously described.^2^ Cryo-electron micrographs were obtained by using core instruments via the Interdisciplinary Center for Biotechnology Research at UF.

### 2D culture

500,000 tumor cells per well in 1-2mL of corresponding culture medium were plated in a 6 well plate overnight to allow plastic cell attachment. Next day, RNA-LPs (5 μg of RNA) in 45μL were added to cells in 2mL of culture media overnight. The following day, microscopy to determine GFP signal was performed as well as collection of supernatant (200μL) was done for downstream evaluation of multiplex chemokine/cytokine evaluation (Eve Technologies, Canada).

### 3D culture

Cells were plated in 3D Darcy plates at initial density of 1×10^6 cell/mL to allow them to form spheroids in LLS polyacrylamide microgel as previously described.^10^ After about 72 hours, spheroid formation was confirmed and viability evaluated. Next, 1mL of tumor spheroid suspension in LLS was collected for each condition and mixed with 45μL of RNA-LP (5μg of RNA). LP alone and media alone were used as controls. Subsequently, 100μL of this mix were plated in 96 well plate with flat glass bottom to allow microscopic evaluation. Each sample was supplemented with 100 μL of corresponding media without disturbing the spheroid-LLS bed. After 24 hours, supernatants were collected for downstream evaluation in a chemokine/cytokine multiplex (Eve technologies, Canada). Tumor spheroids RNA uptake was evaluated by measuring GFP expression through confocal microscope (Nikon A1R HD25).

### In vivo murine experiments

To measure IFN-α2 secretion in serum, wildtype C57BL/6 or MYD88 knock-out mice were intravenously injected with B16F10 tumor cells (300,000 cells) on day 0. On day 1, OVA mRNA-LPs (25 μg RNA in 200 μL) were intravenously injected. Six hours after RNA-LP injection, blood was collected and serum tested for chemokine/cytokine levels. To evaluate effect of checkpoint inhibitors, monoclonal antibodies against murine PD-1 were purchased from BioXcell and given as intraperitoneal loading dose on first day of administration and every 3-4 d thereafter to wildtype C57Bl/6 mice bearing intracranial cortical GL261 or K-luc tumors.

